# Pancreatic changes in obese mice supplemented with vitamin D

**DOI:** 10.1101/2024.11.07.622429

**Authors:** Guilherme da Rocha, Natália Pereira dos Santos, Juliana Cristina Mendonça, Nathan Oscar Baptista Fassis, Bruno Calsa, Marciane Milanski, Maria Esméria Corezola do Amaral

**Author notes:** Corresponding author: Maria Esméria Corezola do Amaral, NUCISA- Common Core of Health, Centro Universitário da Fundação Herminio Ometto, FHO, 13607-339, Araras, SP, Brazil. Co-first authors: Guilherme da Rocha, Natália Pereira dos Santos, Juliana Cristina Mendonça.

## Abstract

This study investigates the impact of vitamin D supplementation on insulin secretion and pancreatic islet morphometry in a high-fat diet-induced obese mouse model. Male C57BL/6 mice were randomly assigned to two groups: a control group receiving a standard diet (C) and an obese group receiving a high-fat diet (H). After 12 weeks, the obese group was subdivided, with one subset continuing on the high-fat diet (H) and the other receiving vitamin D supplementation (HD) at 500 IU/kg via oral gavage for four weeks. Glucose tolerance tests, insulin sensitivity assessments, insulin secretion assays, and histological and immunohistochemical analyses for glucagon and PCNA were conducted. Vitamin D supplementation led to a reduction in fasting blood glucose levels and a significant increase in insulin secretion at 20 mM glucose, as indicated by an improved HOMA%B index, suggesting enhanced β-cell protection. Additionally, vitamin D treatment resulted in a β-cell percentage comparable to the control group and a notable increase in α-cell population relative to the untreated obese group. However, elevated circulating cholesterol levels were observed in both obese groups, with a significant rise in hepatic triglycerides in the vitamin D-treated group. These findings highlight the potential of vitamin D to improve pancreatic secretory function in obese mice.

## Introduction

Studies report that vitamin D has important function in the human body beyond its classic role in bone physiology by regulating calcium and phosphate concentrations. Vitamin D is also known for its involvement in various biological process, such as proliferation and differentiation in different cell lineages, including lymphocytes, endothelial cells and keratinocytes. (National Institutes of Health n.d.). Furthermore, the use of vitamin D participates in modulating diseases such as diabetes, cancer, cardiovascular diseases and inflammation (Mitri et al. 2014; Pfotenhauer and Shubrook 2017).

The reduction in vitamin D concentration is not solely due to low sunlight exposure, it has also been observed in obese individuals, which is linked to body fat accumulation. Evidence suggests that low vitamin D concentration is related to its accumulation in adipocytes, which leads to increased hunger and decreased energy expenditure and leads to insulin resistance and high blood pressure (Didriksen et al. 2015; Konijeti et al. 2016; Zakharova et al. 2019).

Insulin resistance and its relationship with hypovitaminosis D in obese children and adolescents have been reported through measurements of fasting glycemia and serum insulin, as well as homeostatic model assessment (HOMA-IR) (Zakharova et al. 2019).

Studies indicates that the obese pediatric population does not respond to vitamin D supplementation at usual doses as effectively as normal-weight controls (Harel et al. 2011; Aguirre Castaneda et al. 2012). Although there is no consensus on the universally recommended dose for the treatment of hypovitaminosis D in this population, some authors suggest increasing the standard dose (Vidailhet et al. 2012).

Increased levels of glucose and fatty acids in circulation, generally associated with decreased pancreatic function and reduced insulin secretion, are often indicative of insulin resistance and obesity (Purrello and Rabuazzo 2000).

Pancreatic islets from type 1 diabetic donors incubated with vitamin D have shown an increase in vitamin D receptor expression and a reduction in apoptosis (Riachy et al. 2006). Additionally, in vitro study suggested that vitamin D receptor ligand such as calcipotriol (a derivate of 1,25(OH)2D3), may protect human pancreatic β-like cells - derived from pluripotent stem cells - from interleukin-1β-mediated inflammation (Wei et al. 2018). Collectively, these findings indicate that vitamin D can modulate pancreatic insulin secretion through mechanisms involving the renin-angiotensin system (RAS) pathway and may offer protectiong to β-cells against local inflammation process (Folch et al. 1957).

Nevertheless, the precise mechanisms by which vitamin D influences insulin secretion and glycemic control in human pancreatic islets remain to be fully elucidated. In this study, we aim to investigate the biochemical and physiological mechanisms underlying the effects of vitamin D supplementation on pancreatic islets in an obese animal model.

## Material e methods

### Animals

The project received approval from the Animal Ethics Committee under protocol number 021/2021. A total of 30 male C57BL6 mice (*Mus musculus*), 2 months old and with an average weight of 20 grams, were used. The animals were sourced from the Animal Experimentation Center “Prof. Dr. Luiz Edmundo de Magalhães” at the Hermínio Ometto Foundation University Center. They were housed in cages with 4 to 5 mice per cage, maintained at a constant temperature of 23±2°C and 55% humidity, under a 12-hour light/dark cycle, with free access to potable water.

The mice were divided into three experimental groups as follows:

- Control group: Mice fed standard rodent chow ad libitum (n=10) for 4 months.
- Obese group: Mice fed a high-fat diet ad libitum (n=10) for 4 months.
- Obese + Vitamin D group: Mice fed a high-fat diet for 4 months, with oral supplementation of Vitamin D (500 IU/kg/daily) administered during the final month (n=10).

### High-Fat Diet Composition

Containing 60% fat: sucrose 100 g, maltodextrin 155 g, starch 107 g, casein 188 g, lard 310 g, soybean oil 40 g, mineral mix 35 g, vitamin mix 10 g, fiber (cellulose) 50 g, choline bitartrate 2.5 g, L-cystine 2.4 g, for a total of 1000 g.

### Intraperitoneal glucose tolerance test (ip.GTT) and intraperitoneal insulin tolerance test (ip.ITT)

The animals were subjected to Ip.GTT and Ip.ITT after a 6 h fast. Blood was collected from the tail of each rat (time zero). Subsequently, glucose (2 g/kg) or insulin (1.5 U/kg) were administered intraperitoneally, and blood samples were collected at each time point. The glucose concentrations in the blood samples were determined using a glucometer (OptiumXceed, Abbott). Kitt was calculated from the slope of the regression line obtained with log-transformed glucose values between 0 and 30 min after insulin administration. Homeostasis model assessment of insulin resistance (HOMA-IR), homeostasis model assessment of insulin sensitivity (HOMA%S), and homeostasis mode assessment of β-cell function (HOMA-β) were calculated using the website www.dtu.ox.ac.uk/homacalculator.

### Biochemical assessments

After the trial, the animals were euthanized via anesthesia with xylazine (10 mg/kg) and ketamine (90 mg/kg). Biochemical analyses of glucose, triglycerides, cholesterol were performed using commercial kits (Laborlab, Guarulhos, SP, Brazil). Total liver lipids were extracted according to the method of Folch et al. (1957).

### Secretion by isolated islets

Islets were isolated manually after collagenase digestion of the pancreas. Groups of five islets were first incubated for 30 min at 37 °C in Krebs-bicarbonate solution containing 5.6 mmol/L glucose and equilibrated with 95% O 2 /5% CO 2, pH 7.4. The solution was then replaced with fresh Krebs-bicarbonate buffer, and the islets were incubated for an additional hour in the presence of glucose (2 and 20 mmol/L). The incubation medium contained: 115 mmol/L NaCl, 5 mmol/L KCl, 10 mmol/L NaHCO3,

2.56 mmol/L CaCl 2, 1mmol/L MgCl2, 15 mmol/L HEPES, and 0.3% BSA (w/v). The insulin content of each sample was measured using ELISA kit (Biotec Center, SPI-Bio). As a result, the fold increase in insulin release was calculated insulin secretion compared with the basal value observed in the presence of 2 mmol/l glucose when exposed to 20 mmol/l glucose.

### Histological analysis of pancreas

After fixation and dehydration of the pancreas, small fragments were embedded in paraffin using standard procedures. Sections (5 μm thick) were stained with hematoxylin-eosin and Perls (qualitative histological evaluations for Perls were performed by two observers, GR and JCM, considering presence and absence of labeling for hemosiderin - potassium ferrocyanide solution) solution and images were acquired using a Leica DM 2000 photomicroscope and LAS v.4.1 software. For each animal, 10 images were obtained at 400× magnification (1032×1040 pixels). ImageJ software was used to quantify the islet area in cm^2^.

### Immunohistochemistry

For the immunohistochemical study, antigens were recovered from 5 μm thick sections using sodium citrate buffer (10 mM, pH 6.0) at 100 °C for 40 min. Endogenous peroxidases and non-specific binding were blocked using a Novolink Max Polymer Detection System (RE7280-K; Leica Biosystems®). The samples were incubated with the primary antibody PCNA and glucagon (PCNA mouse monoclonal 1:200; Santa Cruz Biotechnology®; anti-glucagon mouse monoclonal 1:1000, Affinity®) overnight. Subsequently, the tissues were incubated with Novolink Polymer (Leica Biosystems), the reaction was developed with diaminobenzidine, and the sections were counterstained with Harris hematoxylin. Five images from each animal at 400× magnification was captured using a Leica DM2000 microscope, and quantified using ImageJ v.1.53e (NIH, USA).

### Statistical analysis

Data are shown as the mean ± standard error of the mean. All data were analyzed using the Shapiro-Wilk Normality test. For comparison between the different experimental groups, one-way ANOVA and Tukey post-test were used. All statistical and graphical tests were performed using GraphPad Prism software (version 8.0). The pre-established significance level was set at p < 0.05.

## Results

### Physical and histology parameters

The body weights obtained for the different animal groups are shown in Figure 1A. The animals of the H and HD groups presented significantly higher weight gain between the 8^th^ and 20^th^ weeks, compared to the C group, which could also be explained by high fat diet. The fat pad index (perirenal adipose tissue/body weight) were greater for the animals of group H and HD groups compared to C group (Figure 1B). The adipocytes areas were higher for group H and HD groups compared to C group (figure 1C). Figure 1D show the adipocyte histology stained with hematoxylin-eosin.

**Figure 1:**
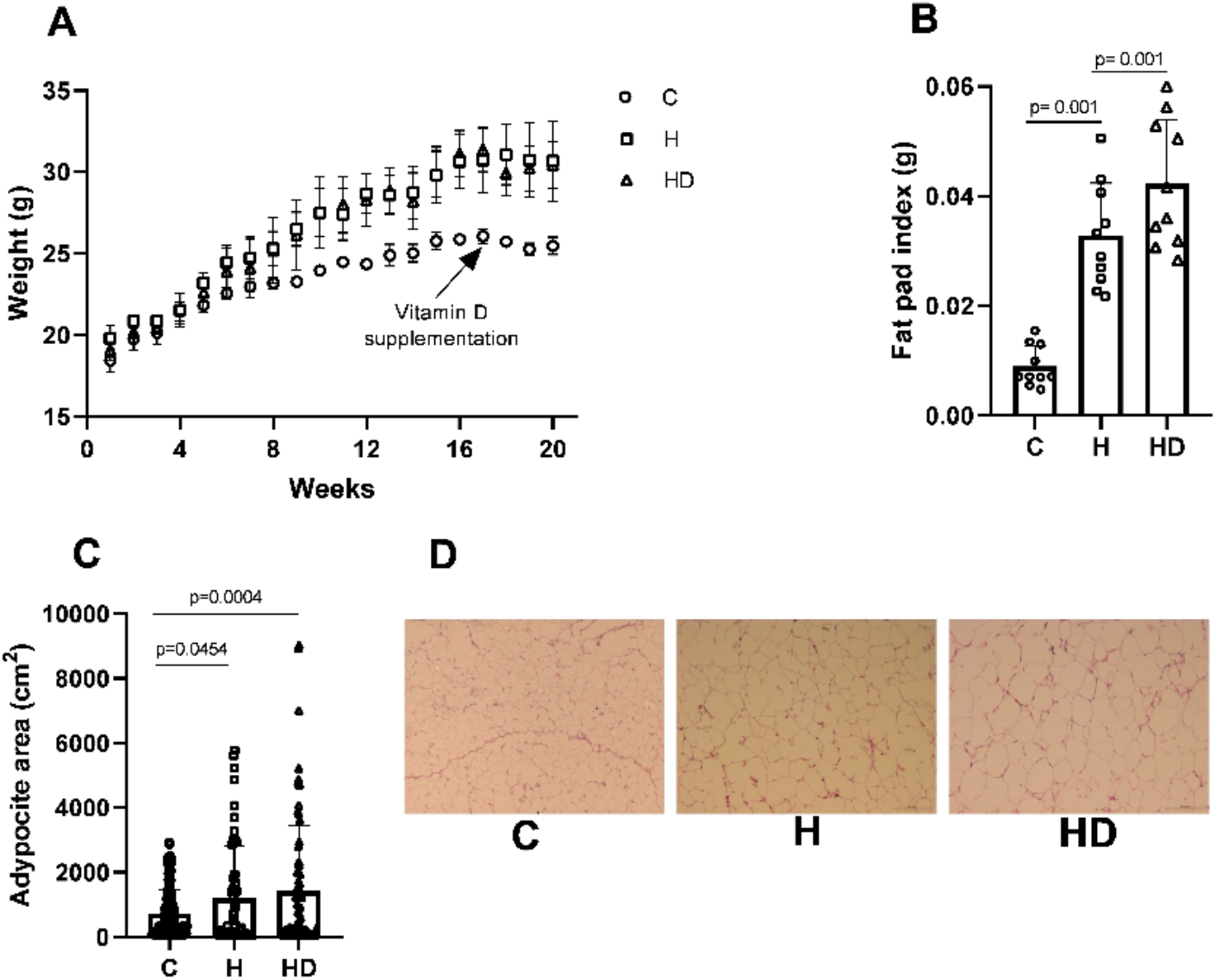
Weight (A), fat pad index (B), adipocyte area (C), morphological adipocyte in H/E (D). Data are expressed by mean ± SEM and were analyzed by one-way ANOVA followed by post-hoc Tukey test. The numbers of mice (n) and p-values are indicated on each graph.

### Fasting glycemia recovery and increased protection of pancreatic islet β cells in animals supplemented with vitamin D

To evaluate the interference of vitamin D in glycemic homeostasis in animal groups, the fasting glycemia test (Figure 2A), HOMA index (Figure 2B, C, D) and glucose tolerance test (Figure 2E, F) were performed. and insulin sensitivity testing (Figure 2 G, H). A reduction in fasting glycemia was observed in animals in the HD group versus H (Figure 2A) and an increase in the HOMA %B index for animals in the HD group compared to animals in groups H and C (Figure 2B). The HOMA%S (Figure 2C) and HOMA IR (Figure 2D) indices present similar values for all animal groups. The glucose tolerance test curve shows characteristics described in the literature for obese animals such as reduced glucose tolerance in the H and HD groups (Figure 2E). This same result was reflected in the area during the glucose tolerance test indicating glucose intolerance in obese, H and HD animals (Figure 2F). The insulin sensitivity test (Figure 2G) and the glucose decay rate analysis (Figure 2H) show similar results between the groups.

**Figure 2:**
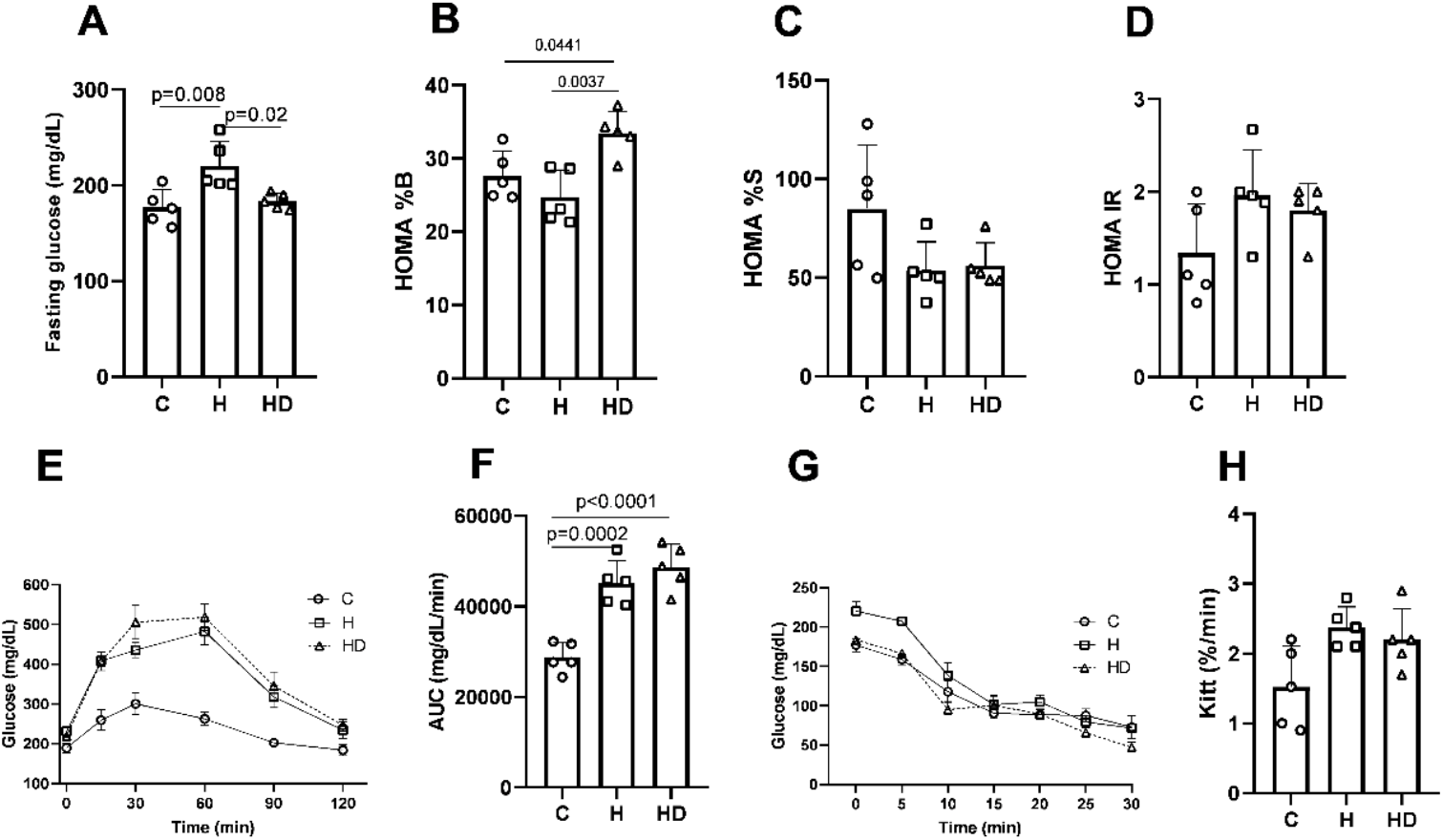
Fasting glucose (A), HOMA %B (B), HOMA %S (C), HOMA IR (D), intraperitoneal glucose tolerance test - ipGTT (E), Area under the curve (AUC) of ipGTT (F), intraperitoneal insulin tolerance test – ipITT (G), rate constant for glucose decay (H). Data are expressed by mean ± SEM and were analyzed by one-way ANOVA followed by post-hoc Tukey test. The numbers of mices (n) and p-values are indicated on each graph.

### Lipid profile change in animals supplemented with vitamin D

There was no change in body weight when comparing the H and HD groups throughout the treatment (data not shown). The concentration of triacylglycerol was similar for all animal groups (Figure 3A). An increase in total circulating cholesterol was observed in groups H and HD compared to group C (Figure 3B) and no difference in VLDL concentrations (Figure 3C). The concentrations of muscle glycogen (Figure 3D) and liver glycogen (Figure 3E) were similar for all groups. An increase in hepatic triacylglycerol was found in animals in the HD group compared to animals in group C (Figure 3F) and hepatic cholesterol concentrations were similar for all groups (Figure 3G).

**Figure 3:**
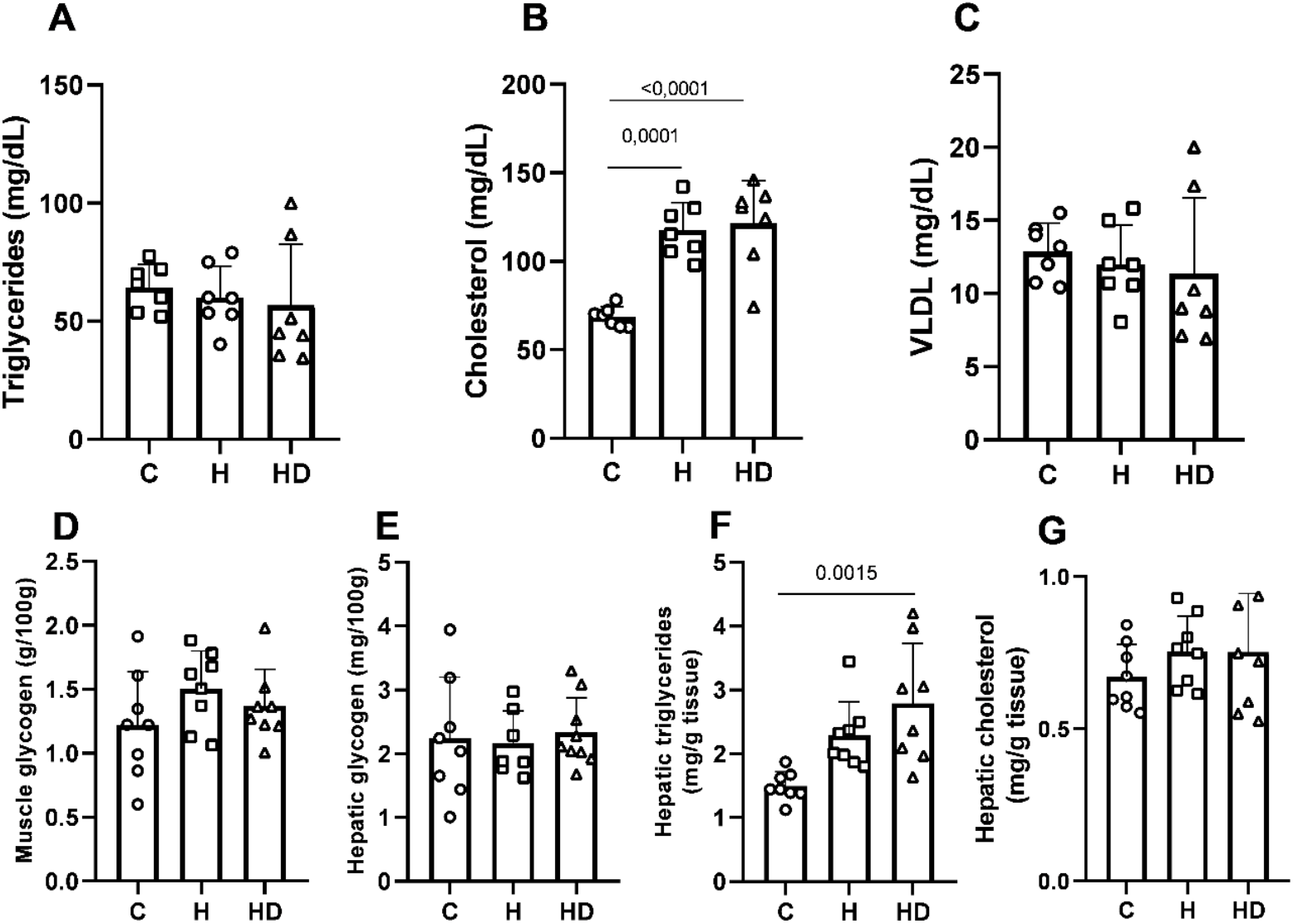
Serum triglycerides (A), serum cholesterol (B), serum very low density liprotein (C), muscle glycogen (D), hepatic glycogen (E), hepatic triglycerides (F), hepatic cholesterol (G). Data are expressed by mean ± SEM and were analyzed by one-way ANOVA followed by post-hoc Tukey test. The numbers of mices (n) and p-values are indicated on each graph.

### Insulin profile, glucose-stimulated insulin secretion and immunohistochemical evaluation for PCNA and glucagon in pancreatic islets

Fasting insulinemia was similar for all animal groups (Figure 4A). Insulin secretion by isolated islets stimulated in 2mM glucose was also similar for the groups (Figure 4B). However, insulin secretion by isolated islets stimulated by 20mM glucose (Figure 4C) and increased insulin secretion at 20mM (Figure 4D) were greater for animals in the HD group compared to animal groups C and H. The area of the islet of animals in group H was larger when compared to animals in group C and HD (Figure 4E). The count of PCNA antibody-positive cells in pancreatic islets was similar for all animal groups (Figure 4F). The percentage of β cells (Figure 4G) and the percentage of α cells (Figure 4H) were higher in animals supplemented with vitamin D compared to animals in groups C and H. Figure 4I shows the representative plate of the islets from the analyzes histological and immunohistochemical analyses.

**Figure 4:**
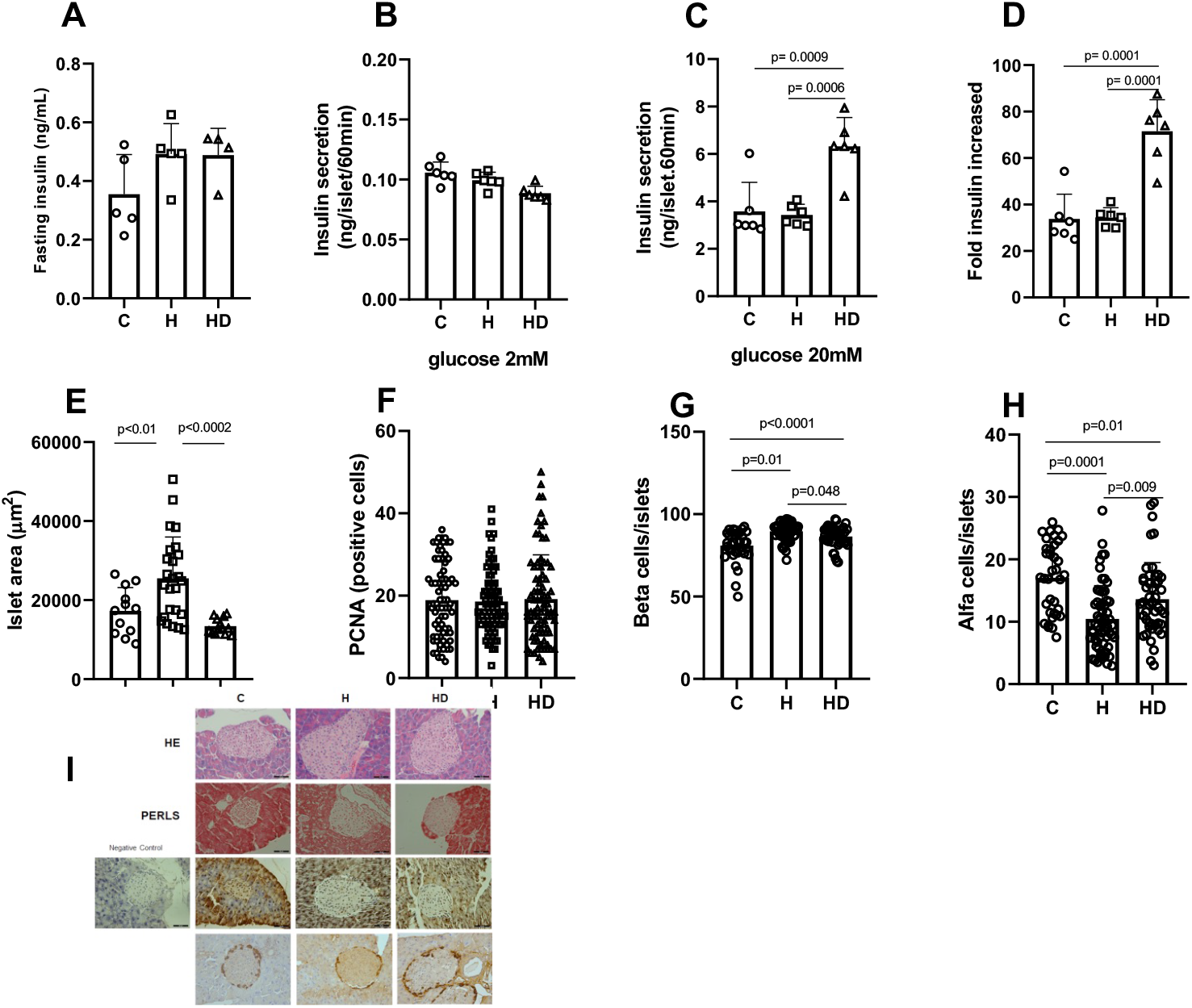
Fasting insulin (A), insulin secretion under 2 mM glucose (B), insulin secretion under 20 mM glucose (C), fold insulin increase (D), islet area (E), PCNA immunostaining (F), percentage of beta cells (G), percentage of alfa cells (H), morphological and immunostaining assessments in islet (I). Data are expressed by mean ± SEM and were analyzed by one-way ANOVA followed by post-hoc Tukey test. The numbers of mice (n) and p-values are indicated on each graph.

## Discussion

In the present study, the benefits of vitamin D in reducing fasting glycemia were evidenced, as reflected by the HOMA%B index, the increased in insulin secretion stimulated by 20 mM glucose and histomorphology characteristics of the islets.

The relationship between reduced vitamin D and both type 1 and 2 diabetes has been reported by several researchers. (Littorin et al. 2006; FENG et al. 2017; Hafez et al. 2017; Rasouli et al. 2022)

Our findings are consistent with studies on islets isolated from vitamin D receptor knockout (VDR-KO) mice (Cheng et al. 2011) and with studies involving mice with a mutation in the AF2 domain of the vitamin D receptor, which prevents ligand-induced transcriptional activation of vitamin D-responsive genes. (Vangoitsenhoven et al. 2016) However, in contrast to results obtained from mice that functionally inactivated vitamin D receptor (Vangoitsenhoven et al. 2016), we found no difference in glucose tolerance or insulin sensitivity evidenced by GTT and ITT, respectively, in animals supplemented with vitamin D (HD) compared to hyperlipidic animals (H).

Studies involving VDR-KO mice show decreased insulin secretory capacity and glycemic control (Cheng et al. 2011). Furthermore, the same mutant mouse (Zeitz et al. 2003) showed a reduction in maximum insulin secretion. Transgenic mice that overexpressed the vitamin D receptor in pancreatic β-cells were protected from streptozotocin-induced diabetes and showed preservation of β-cell mass and reduced inflammation, reinforcing our findings. (Morró et al. 2020)

Vitamin D increases the expression of the insulin gene and may be involved in organization of the cytoskeleton and cellular growth of beta cells. (Wolden-Kirk et al. 2013; Altieri et al. 2017) Additionally, vitamin D is involved in the flow of calcium, depolarization of pancreatic beta cells and insulin release. (Alvarez and Ashraf 2010) Our data agree with these studies and contribute to the elucidation of the effects of vitamin D in cases of hyperglycemia and obesity.

The therapeutic role of vitamin D in lipid abnormalities has been demonstrated. (McGill et al. 2008; Asemi et al. 2013) Vitamin D at a dose of 500 IU/Kg orally for 5 weeks modified lipid abnormalities in obese rodents by reducing serum concentrations of LDL, triacylglycerol and total cholesterol. Results from our work demonstrate that daily oral dose of 500U/Kg did not modulate lipid profile of obese animals supplemented with vitamin D. Instead, we observed an increase in triacylglycerol in the liver of these animals.

GC-globulin (GC), or vitamin D-binding protein, is a multifunctional protein involved in the transport of vitamin D. In pancreatic islets, the gene encoding GC is located in glucagon-secreting α cells. GC contributes to the maintenance of α-cell function. (Viloria et al. 2020, 2022) Deletion of GC results in smaller, hyperplastic α cells that exhibit abnormal Na+ conductance, Ca2+ fluxes, and glucagon secretion. (Viloria et al. 2023) These data from the literature may corroborate our findings on the increase in glucagon-secreting α cells.

Finally, the dose discrepancy, treatment duration, differences between human and animal models underscore the need for further research into the effects of vitamin D to develop new therapeutic approaches for the treatment of obesity.

## Statements and Declarations

## Acknowledgments

We thank Ana Cristina Pires Menegheti, Bruno Alves Cia, Mateus Eduardo Bortolanza da Silva, Renata Barbiere and Letícia Franco for technical assistance.

